# A duo of fungi and complex and dynamic bacterial community networks contribute to shape the *Ascophyllum nodosum* holobiont

**DOI:** 10.1101/2025.03.20.643298

**Authors:** Coralie Rousseau, Gwenn Tanguy, Erwan Legeay, Samuel Blanquart, Arnaud Belcour, Sylvie Rousvoal, Philippe Potin, Catherine Leblanc, Simon M. Dittami

**Affiliations:** Integrative Biology of Marine Models (LBI2M, UMR 8227), Station Biologique de Roscoff, Sorbonne University, CNRS, 29680 Roscoff, France; Genomer Platform, FR2424, Station Biologique de Roscoff, Sorbonne University, CNRS, 29680 Roscoff, France; Genomer Platform, FR2424, Station Biologique de Roscoff, Sorbonne University, CNRS, 29680 Roscoff, France & Adaptation and Diversity in the Marine Environment (UMR 7144), Station Biologique de Roscoff, Sorbonne University, CNRS, 29680 Roscoff, France; Dyliss, Inria (UMR 6074), CNRS, University of Rennes, Rennes, France; MICROCOSME, Inria, 38100 Grenoble, France

**Keywords:** Macroalgae, *Ascophyllum nodosum*, metabarcoding, bacteria, fungi, co-occurrence network, host normalization

## Abstract

The brown alga *Ascophyllum nodosum* and its microbiota form a dynamic functional entity named holobiont. Some microbial partners may play a role in seaweed health through bioactive compounds crucial for normal morphology, development, and physiological acclimation. However, the full spectrum of the microbial diversity and its variations according to algal life stage, season, and location have not been comprehensively studied. This study uses 208 short-read metabarcoding samples to characterize the bacterial, archaeal, and microeukaryotic communities of *A. nodosum* across three nearby sites, four thallus parts, and a monthly survey, aiming to explore the dynamics of ecological interactions within the holobiont. Our results revealed that *A. nodosum* harbors a predominantly bacterial microbiota, varying significantly across all covariables, while archaea were virtually absent. An innovative normalization using the co-amplified host reads provided an estimation of bacterial abundance, revealing a drastic decline in May, potentially linked to epidermal shedding. In contrast, fungal communities were stable, dominated by *Mycophycias ascophylli* and *Moheitospora* sp., which remained closely associated with the host year-round. We identified a core microbiome of 22 ASVs, consistently found in all samples, including *Granulosicoccus,* a genus consistently abundant in other brown algal microbiota. Sequence clustering revealed multiple species which vary according seasons, even in the overall stable *Granulosicoccus* genus. Co-occurrence network analysis revealed putative interactions between microbial groups in response to ecological niches. Overall, these findings highlight the dynamic of bacterial interactions and stable fungal associations within the *A. nodosum* holobiont, providing new insights into the ecology of its microbiota.

## Introduction

Brown seaweeds provide habitats for diverse communities of epi- and endosymbiotic microorganisms (i.e., bacteria, eukaryotes, archaea, and viruses) [1, 2]. The algal surface is suitable for the development of microorganisms because of organic substances, which play an important nutritional role for the microbiota [3]. Some microbial partners are, in turn, necessary for seaweed health by producing bioactive compounds essential for normal morphology, development, and physiological responses to environmental factors [4]. These host-microorganism associations constitute interacting and sometimes co-dependent units, so-called holobionts [2, 5–7].

*Ascophyllum nodosum* is abundant and perennial along the North Atlantic coasts, anchoring to rocky substrates in sheltered intertidal zones. It can live up to 15 years, reach lengths of up to 2 meters, and exhibits slow growth through apical elongation, forming air bladders annually [8–10]. This dioecious species has a slow recruitment process due to limited dispersal and low germling survival rates [11, 12], with young individuals requiring approximately five years to reach maturity [13]. *Ascophyllum* is a valuable resource due to its bioactive components used in agriculture, food ingredients, health, and cosmetics. However, increasing demand has raised concerns about overexploitation.

*A. nodosum* has been described as engaged in a tripartite symbiosis with the endophytic marine fungus *Mycophycias* (formerly *Mycosphaerella*) *ascophylli* and the epiphytic red alga *Vertebrata* (formerly *Polysiphonia*) *lanosa*. The ascomycete *M. ascophylli* is consistently present throughout the host, without apparent pathogenic effects [14, 15], and is thought to protect *A. nodosum* germlings from desiccation [16]. *V. lanosa* is an obligate epiphyte, largely restricted to *A. nodosum,* causing limited damage to the host despite rhizoid penetration into host tissues [17, 18]. These species have undergone considerable coevolution resulting in intricate morphological, physiological, and ecological interactions. However, it is unclear whether these interactions also involve other bacteria or fungi, and whether these fungi and bacteria can increase *A. nodosum* fitness, as described in other brown algae [19].

*A. nodosum*-associated bacteria were first described in 1969 by culturing [20] and later by electron microscopy [21]. In 1988, experiments suggest that bacteria shield *A. nodosum* from digestion by *M. ascophylli* [22]. Further studies investigated surface-attached bacteria degrading polysaccharides [23], the role of endobacteria in quorum-sensing [24], and epibacterial community responses to stress [25] and decay [26]. Recently, Brunet *et al.* [27] quantified the epibacterial communities by qPCR across a year-long survey. Despite recent increased scientific interest, the full diversity of microbial communities (bacteria, eukaryotes, archaea, and viruses) associated with *A. nodosum* remains largely unexplored.

This study aims to fill the knowledge gap across three key questions: (i) Does the microbiota change across a small spatial scale, algal thallus parts, and throughout a monthly survey? (ii) Is there a core microbiota? (iii) How do microorganisms associate with each other? To comprehensively describe the holobiont, we used a short-read metabarcoding analysis of bacteria, archaea, and eukaryotes and used co-occurrence networks to infer potential ecological interactions between microorganisms.

## Material and methods

### 1. Sample collection

A monthly survey was conducted from November 2021 to November 2022 at Pleubian (48.849411, −3.078291, Brittany, France, see map [28]), a location known for the commercial harvesting of seaweeds, including *A. nodosum*. The microbial community associated with this alga was characterized in three nearby sampling sites. At each site, four algae were selected arbitrarily (see [28]) and four parts were sampled: apex, receptacle (during reproductive season only), medium, and basal parts. Algal samples (∼50 mm long) were cut with clean scissors rinsed with 70% ethanol, placed in cryotubes, and stored on ice. In the laboratory, samples were stored at −80°C.

### 2. DNA extraction and sequencing

The algal samples were free-dried and finely ground to access the epi- and endophytic microbial communities. The DNA extraction was based on CTAB extraction buffer, chloroform/isoamyl separation, and the extracted DNA was purified twice with a NucleoSpin Plant II kit (MachereyNagel, Germany) and with AMPure XP Beads (Beckman Coulter, Brea, CA, USA) [28]. To reduce the number of samples and sequencing costs, and facilitate the bioinformatics analysis, three amplicon datasets were built based on the following research questions by pooling at equal concentration the DNA:

- “Site”: Does microbiota vary between the three sampling sites? The strategy involved pooling extracted DNA from the three thallus parts of one individual to remove the potential thallus part effect (apex, medium and base – no receptacle as it was not always present). Four pooled DNA samples from each site were examined for two arbitrarily chosen sampling dates: November 2021 and March 2022 (3 sites * 4 individuals * 2 months = 24 samples)
- “Algal parts”: Does the microbiota vary depending on the part of the alga? Samples from four individuals across four thallus parts were analyzed from site 1 for two arbitrarily choses sampling dates: January and April 2022 (4 parts * 4 individuals * 2 months = 32 samples).
- “Season”: Does the microbiota change with the season? The strategy involved pooling extracted DNA from three thallus parts from one individual alga collected over 12 months at site 1 (12 months * 4 individuals = 48 samples).

We furthermore included three replicates each of a bacterial and fungal mock community containing 10 bacterial and 10 fungal DNAs [28], as well as 8 negative controls. PCR reactions and library preparation were carried out as described in [28], using the universal primers 515-F/926-R [29] to target bacteria, archaea, and eukaryotes, and ITS2 nested PCR (ITS-1/ITS4 and 5,8S-FUN/ITS4-FUN, [30, 31]) to increase taxonomic resolution for fungi. Libraries were sequenced on a NovaSeq 6000 SP at the Genomics Core Facility GenoA platform (Nantes, France) using 500 cycles (kit ref. 20028402).

### 3. Bioinformatic analysis

#### 2.1 Cleaning, taxonomy assignment, and decontamination steps

Bioinformatics analyses were carried out as described in [28]. Briefly, the 16S and 18S (i.e., SSU) and ITS sequences were quality-controlled, and primers were removed with Cutadapt [32]. For the SSU sequences, the analysis was carried out by following the online pipeline (https://astrobiomike.github.io/amplicon/16S_and_18S_mixed) to separate 16S and 18S, with magicblast tool [33]. Each set of 16S, 18S, and ITS files was filtered independently where the forward and reverse reads of the 16S and ITS datasets were merged, and those of the 18S dataset were concatenated. They were denoised into Amplicon Sequence Variants (ASVs) with the DADA2 v1.28.0 [34] in R v.4.2.3. Taxonomy was assigned using SILVA NR99 v138.1, PR2 v.5.0.0, and UNITE v.9, respectively. Short unassigned ITS sequences were compared with full-length ITS consensus sequences from our previous study [28] to aid their assignment. SSU sequences shorter than <350 bp and plastid sequences were deleted. The remaining sequences were decontaminated using microDecon package [35]. All sequences containing homopolymers longer than 5 bases were also excluded. Finally, remaining chimera were removed using vsearch v.2.22.1 [36] with the *--uchime_ref* option and the SILVA non-redundant small subunit database v.138.1 as reference.

#### 2.2 Dataset normalizations

As the universal primer amplified also the 18S of the *Ascophyllum nodosum* host, we were able to use these host reads as an internal reference to detect variations in the proportion of microbial reads: microbial read abundances were divided by the abundance of *A. nodosum* reads (Supplementary Figure 1) in the same sample and multiplied by the median of all *A. nodosum* reads across all samples. As the Illumina sequencing is insufficient to discriminate sequences among the Fucaceae family, all ASVs identified as “*Fucus*”, “*Silvetia*” and “Phaeophyceae” were summarized as *Ascophyllum* for this purpose.

SSU ASVs accounting for less than 0.005% of the total reads in all samples were removed from datasets after normalization. As the ITS primers did not amplify the host, the fungal dataset was normalized by the median and the rare ASVs with under 0.01% relative abundance were removed for the three datasets. This threshold was increased compared to the SSU dataset considering that initial number of sequenced reads for the ITS was lower.

#### 2.3 Beta and alpha diversities

The beta diversity was estimated from raw datasets using the Bray-Curtis dissimilarity index with non-metric multidimensional scaling (nMDS) ordinations and statistical analyses were based on a PERMANOVA test with 999 permutations. The alpha diversity was further calculated with the Shannon index on normalized data using the microbiome package v1.22.1 and compared using ANOVA tests (stats package v.4.3.3) and the Tukey post-hoc tests (rstatix package v.0.7.2).

We removed the sample L1B4_APRIL as it was an outlier in the NMDS plot and because it contained 4 times fewer reads than the average sample, maybe because of a low initial DNA concentration (2.96 ng/μL). Then, the “Sites”, “Algal parts” and “Season” datasets were processed separately. L1I1_MARCH from the “Sites” and “Season” datasets and L1R3_APRIL from “Algae parts” showed inconsistent behavior among multiple analyses such as alpha and beta diversities and barplots. To confirm if these two samples are outliers, a Grubbs test on read abundances in Past4 v.4.15 [37] was performed and confirmed the hypothesis with a p-value < 0.05. They were removed for the rest of the analyses.

#### 2.4. Core microbiota

Intermediate core microbiota was first identified separately for the “Algal parts” and the “Sites and Season” datasets with Venn diagrams (MicEco package v.0.9.19) and pie charts (ggplot2, package v.3.5.0). The intersection of both intermediate core microbiota then formed the global core community of *A. nodosum*. Only ASVs present in 100% of the samples were considered part of the core.

#### 2.5 Heatmap

To distinguish different fungal ecotypes, especially concerning *Moheitospora* sp. or *Mycophycias ascophylli*, all three ITS datasets were merged and the sequences were clustered at 99% identity (e.g., Operational Taxonomic Unit, OTU) with vsearch v.2.22.1 [36]. OTUs were filtered to remove rare OTUs with under 0.01% relative abundance over the entire dataset. Clustered sequences were aligned with Mafft v.7 using the iterative refinement method G-INS-i. The alignment was curated with Gblocks allowing smaller final blocks, gap positions within the final blocks, and less strict flanking positions and a phylogenetic tree was calculated on phylogeny.fr [38] using PhyML, the GTR+G+I substitution model, and an approximate Likelihood-Ratio test. A heatmap was generated with ggplot2 package v.3.5.0.

#### 2.6 Co-occurrence network

For the co-occurrence network, both bacterial and fungal sequences were clustered at 99% identity as above. They were separately normalized by the median and then merged. The merged dataset was filtered with a minimum occurrence larger than 45 reads in at least 3 samples [39]. Putative associations between covarying OTUs were inferred with SpiecEasi v.1.1.3 [40] using the “glasso” probabilistic inference method and a lambda.min.ratio of 0.01. Significant covariances were converted to correlations with the function cov2cor [39]. Only positive edges were considered because most module detection algorithms allow only positive edges. The Leiden module algorithm detection was chosen after comparing eight algorithms (Fast Greedy, Infomap, Label Propagation, Leading Eigenvector, Leiden, Louvain, Spinglass, and Walktrap) using the igraph R package v.2.0.3 [41] as reported in [42]. To visualize the network, the Gephi software v.0.10.1 [43] with the Force Atlas2 algorithm was used. Metabolic capabilities from the taxonomic affiliations of each module were inferred using EsMeCaTa v.0.5.4 [44]. The metabolic enrichments, implemented in EsMeCaTa, were performed with Geasapy [45] and the resulting heatmap with Orsum [46].

## Results

### 1. Shifts in seasonal bacterial diversity and global community patterns across datasets

The alpha diversity of the *Ascophyllum*-associated microbiome in the SSU dataset varied significantly over the year (Figure 1A) but not between “sites” and “algal parts” (Supplementary Figure 2). The diversity was the lowest in May 2022 and differed significantly from the samples in November 2021, March, June, September, and October 2022 (Figure 1A). Concerning the ITS diversity, no significant differences were detected in any dataset (Supplementary Figure 3).

**Figure 1.**
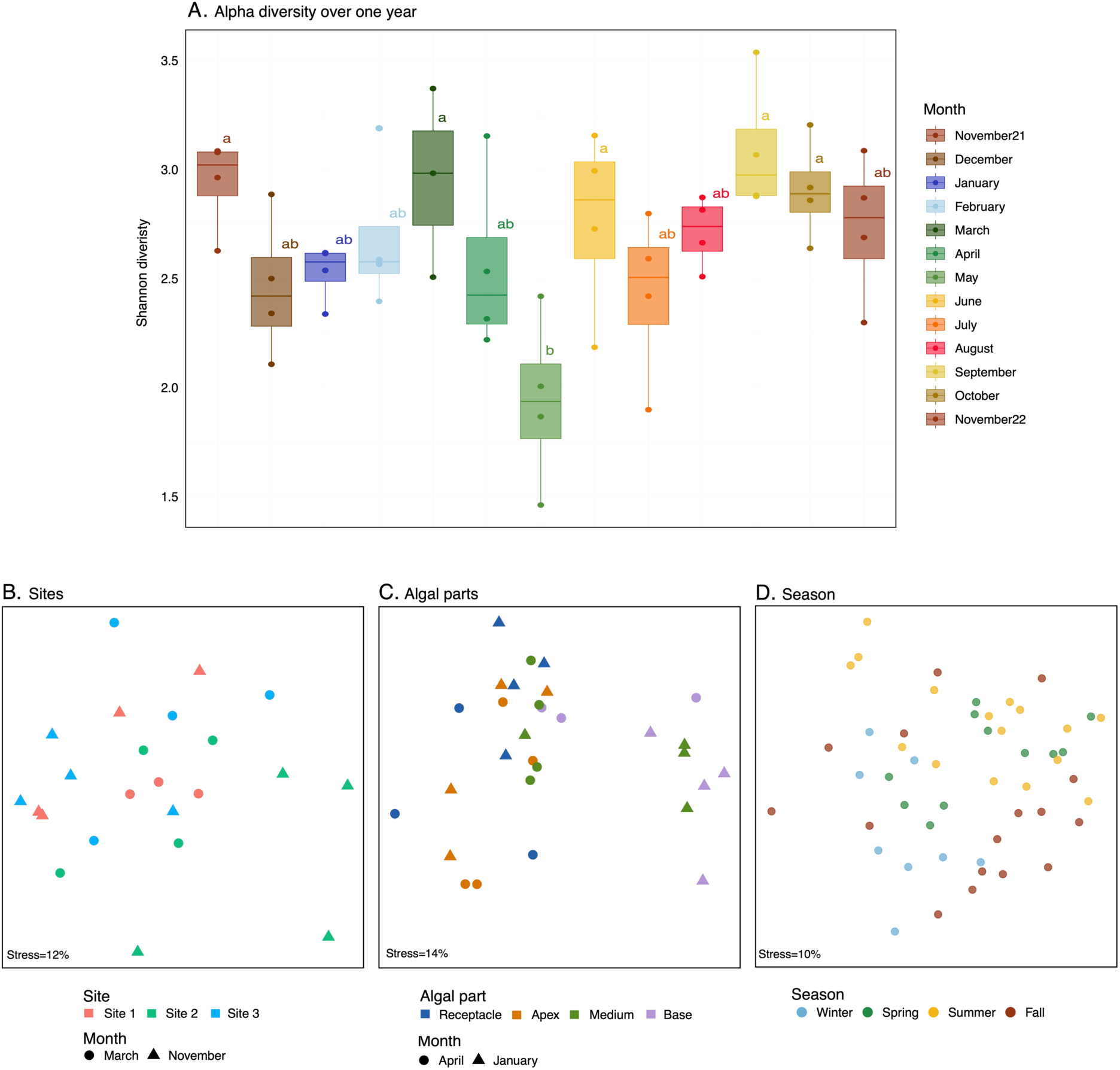
| Variation of SSU diversity and communities across the three datasets. A. Alpha diversity (Shannon index) of season dataset. Significant differences according to an ANOVA with a Tukey post-hoc test are indicated by different letters next to the bars. B. The NMDS plot concerns the beta diversity of the “Sites” dataset and shows significant differences for months (p-value=0.015) but not for sites. C. The NMDS concerns the “Algal parts” dataset and shows significant differences for the algal parts (p-value=0.001) and month (p-value=0.001). D. The NMDS concerns the “Season” dataset and shows significant differences for season (p-value=0.001) and month (p-value=0.001). Statistical analyses were based on a PERMANOVA test.

The SSU communities were stable at a small geographic scale in November 2021 and March 2022 (Figure 1B). Along the algal thallus, different groups emerged, separated by months and thallus parts (Figure 1C). In January, the base and medium parts were similar, except for one base from April, and distinct from the receptacle and apex. Seasonal variations were observed with distinct cluster between fall/winter and spring/summer (Figure 1D). ITS profiling revealed only one significant difference between January and April in the “algal parts” dataset (Supplementary Figure 4).

### 2. Dynamic and highly diverse bacterial communities: dominance of seven classes

The *A. nodosum* bacterial communities are mainly composed of 7 classes (Figure 2), with *Alphaproteobacteria* and *Gammaproteobacteria* consistently dominating. The most abundant genera include an unassigned Alphaproteobacterial genus, *Granulosicoccus* (Gammaproteobacteria), *Pleurocapsa PCC-7319* (Cyanobacteria), *Blastopirellula* (Planctomycetes), and *Algisphaera* (Phycisphaerae).

**Figure 2.**
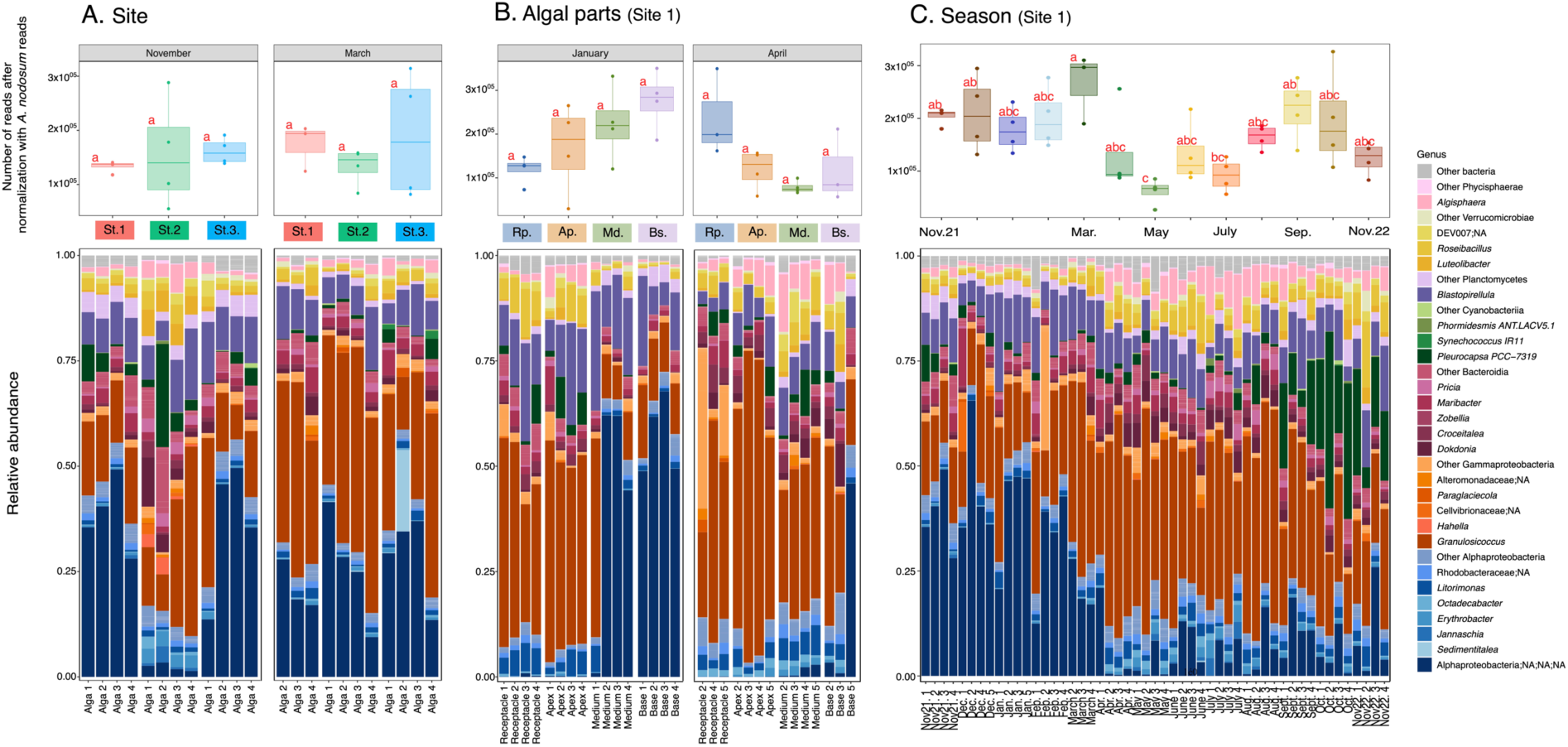
| Variation of the 16S diversities of *Ascophyllum nodosum* at Pleubian sites (Brittany, France). The boxplot figures at the top show the number of total bacterial reads after normalization with *A. nodosum* nuclear 18S rDNA reads (see methods section 2.2). Here significant differences are estimated with an ANOVA and a Tukey post-hoc test and indicated by different letters next to the bars. No letter means no significant differences. The barplots show the variation in relative abundance of bacteria at the genus levels for the “Site” (A. St1=Site 1, St2=Site 2, St3=Site 3), “Algal parts” (B. Rp=Receptacle, Ap=Apex, Md=Medium, Bs=Base) and “Season” (C). The different shades of blue represent the Alphaproteobacteria, orange the Gammaproteobacteria, dark pink Bacteroidia, green Cyanobacteriia, violet Planctomycetes, yellow Verrucomicrobiae, pink Phycisphaerae, and grey other bacteria in small abundance, not included in the 7 major classes. “NA” corresponds to unassigned taxonomy.

#### Site dataset

Bacterial abundances relative to the host remained relatively stable across sites and months (Figure 2A, boxplot) - despite site-dependent variations in terms of data dispersion - except at site 2 in November, where the unassigned Alphaproteobacterial genus was less abundant than in other samples.

#### Algal parts dataset

No significant differences were detected in bacterial read abundance relative to the host, in the different algal parts. Regarding the community structure, *Granulosicoccus* was the dominant genus in every sample except for the medium and basal parts in January where the unassigned Alphaproteobacterial genus were found in high relative abundance (Figure 2B). We also separately analyzed the abundance of host plastid reads relative to nuclear reads across algal parts, revealing a significantly lower proportion of plastid reads in the receptacles (Supplementary figure 5B).

#### Seasonal dataset

Normalized bacterial read abundance was relatively stable from November 2021 to March 2022 (Figure 2C, boxplot) before it drastically dropped in April and May and then increased from July to October. These observations are supported by an ANOVA and a Tukey post-hoc test, which yielded significant results in November 2021, December, March, May, July, and September, with all significantly differing from May.

From November 2021 to March 2022, the bacterial communities were dominated by one ASV affiliated to an unassigned Alphaproteobacteria, but this group was less abundant from April to November 2022. From August, the genus *Pleurocapsa* PCC-7319 increased in abundance to reach the highest levels in September, October, and November 2022. The ratio of plastid to nuclear host reads was also highly variable across samples (up to 5-fold difference), although no significant seasonal pattern was detected (Supplementary figure 5A).

### 4. Stable eukaryotic communities dominated by two fungi

Fungi belonging to Ascomycota class dominated the 18S sequences (>70% of total reads), in particular representatives of the classes Dothideomycetes and Sordariomycetes (Figure 3.1). Similarly, for the ITS sequences two dominant fungi identified as *Mycophycias ascophylli* (Dothideomycetes) and *Moheitospora* sp. (Sordariomycetes, recently renamed *Juncigena* sp.) were observed in almost every sample (Figure 3.2).

**Figure 3.**
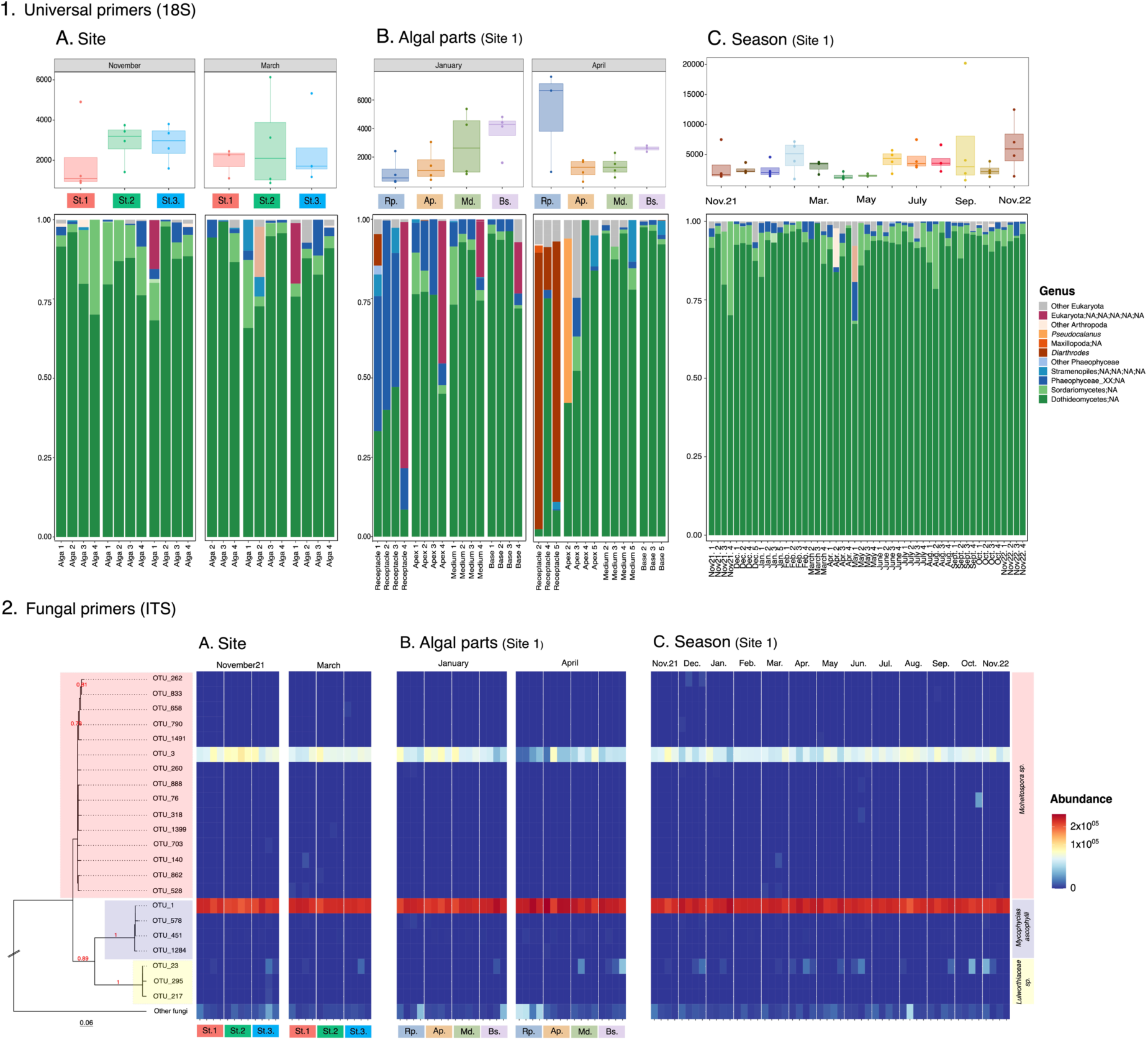
| 18S (1.) and ITS (2.) based-primers diversities of the *Ascophyllum nodosum* microbiome at Pleubian sites (Brittany, France). The boxplot figures at the top show the number of eukaryotic reads after normalization with *A. nodosum* nuclear rDNA reads (see methods section 2.2). The barplots (1.A, 1.B, 1.C) show the variation of eukaryotes in relative abundance at the genus levels for the “Site” (A. St1=Site 1, St2=Site 2, St3=Site 3), “Algal parts” (B. Rp=Receptacle, Ap=Apex, Md=Medium, Bs=Base) and “Season” (C). The different shades of green represent the Ascomycota class, blue the Phaeophyceae, orange the Arthropoda, dark pink non-assigned genera at the phylum level, and grey other eukaryotes found in small abundance, not included in 4 major families. “NA” corresponds to unassigned phylogeny. The heatmap shows the variation of fungal strains across the three datasets (2.A, 2.B, 2.C). ASV were clustered at 99% (OTUs) of identity to distinct strains. OTUs not included in the three abundant species (*Mycophycias ascophylli*, *Moheitospora sp.* and *Lulworthiaceae sp.)* were summed in the “Other fungi” category. The stronger the red, the more abundant is the OTUs. The phylogenetic tree on the left was based on a sequence alignment of 22 sequences with the MAFFT (G-INS-I iterative strategy), curation with Gblocks (329 positions remained), and was calculated using PhML (SH-LIKE, GTR+G+I substitution model). Support values under 0.70 have been removed and there is no support value for the ‘other’ category because it represents several fungi. The three most abundant fungal genera were highlighted in the tree: red for *Moheitospora sp.*, blue for *Mycophycias ascophylli,* and yellow for *Lulworthiaceae sp*.

The host-normalized eukaryotic read abundance remained stable across datasets (Figure 3.1.A, 3.1.B, 3.1.C, boxplots), following similar patterns as the bacterial communities, except that seasonal variations were less pronounced (Figure 3.1.C). In the algal parts dataset (Figure 3.1.B), however, Phaeophyceae_XX;NA were abundant in January in the receptacles and then decreased from apex to basal parts. In April, this group was observed in low abundance, and two of three receptacles were dominated by the *Diathrodes* genus in the Arthropoda phylum.

### 5. The *A. nodosum* core microbiome

The four-thallus parts shared 28 ASVs and a total of 60 ASVs were consistently present only on specific algal parts: 16 ASVs were consistently observed only in receptacles, 4 in apex, 10 in medium, and 30 in basal parts (Figure 4A.1). The core community specific to basal parts was the most taxonomically diverse with 7 different classes, followed by that of the receptacles, the medium and the apical parts, the latter comprising only 3 classes in the core community (Figure 4A.3).

**Figure 4.**
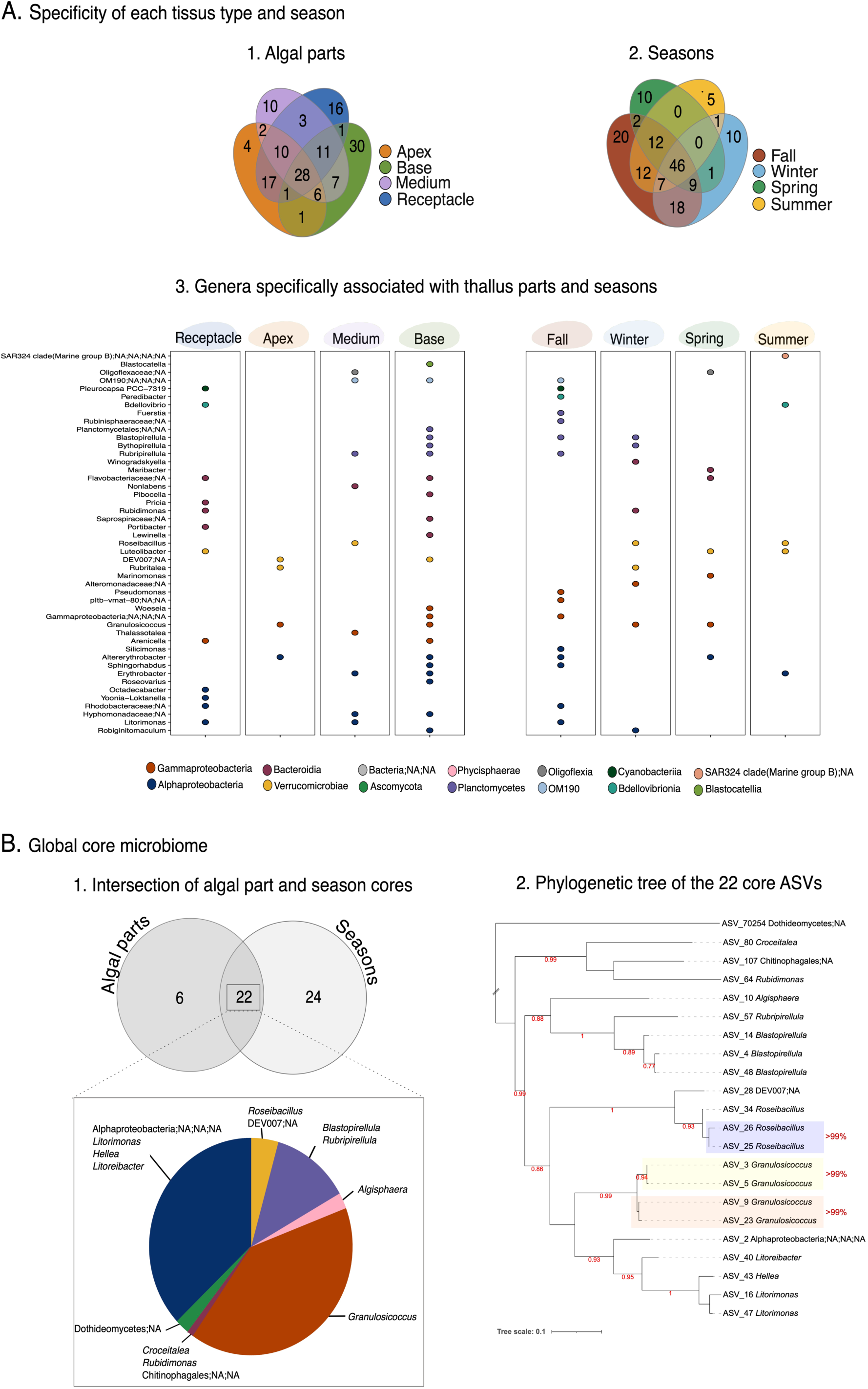
| Core microbiome (Bacteria and Eukaryota, SSU) of *Ascophyllum nodosum* specifically associated with the four algal parts (A.1) and over the four seasons (A.2) (i.e., “intermediate core microbiota”). Numbers in the Venn diagrams represent the number of ASVs. The bubble plot (A.3) represents genera consistently associated specifically with the four algal parts and the four seasons. The global core microbiome is formed by the intersection of both intermediate core microbiota (B.1). The pie chart shows the genera these 22 ASVs belong to and their relative read numbers in the global core microbiome. The color shows the bacterial class. The phylogenetic tree (B.2) was based on a sequence alignment of 22 core sequences with the MAFFT (G-INS-I iterative strategy), curation with Gblocks (373 positions remained), and was calculated using PhML (SH-LIKE, GTR+G+I substitution model). Support values (SH-like as approximate Likelihood-Ratio test) under 0.75 have been removed. The ASV sharing more than 99% of identity, performed with vsearch tool, were highlighted in blue, yellow and orange.

In the seasonal dataset, we identified 46 ASVs present on every individual the entire year, and another 45 ASVs were found consistently only in one season (20 ASVs observed only in fall, 10 in winter, 10 in spring, and 5 in summer; Figure 4A.2). The fall microbiome also exhibited the most diversified core community with 6 classes, followed by winter, spring and summer (Figure 4A.3).

The intersection of the core of the algal parts and season, comprised 22 ASVs with a large dominance of *Granulosicoccus*, followed by an assigned Alphaproteobacteria ASV (Figure 4B.1). Among the 22 ASVs, two pairs of *Granulosicoccus* and *Roseibacillus* shared more than 99% of identity (Figure 4B.2).

### 6. Putative microbial interactions: the co-occurrence network analysis

We investigated groups of co-varying OTUs that could share the same ecological niches via a co-occurrence network analysis. After merging SSU OTUs with ITS OTUs, the final matrix contained 12,090 OTUs. After filtering out low-abundance OTUs, 531 OTUs remained for co-occurrence analyses, representing 99.3% of the total reads in the dataset. A total of 520 OTUs were correlated with 4130 positive (72.6%) and 1552 negative (27.3%) edges. Only 11 OTUs were isolated. For the visualization, we only used the positive correlations.

In the inferred network, 8 modules comprising OTUs with similar distribution patterns were identified (Figure 5A, B). Module 4 was dominant with 88.1% of reads (including *Ascophyllum*, *Mycophycias*, *Moheitospora* and *Granulosicoccus*), followed by seven modules representing between 6.5% and less than 1% of the reads. Among the 19 OTUs found in the core microbiome, 14 OTUs fell into module 4, followed by module 8 with 3 OTUs, and module 1 and 6 (1 OTUs each) (Figure 5A). Some of the core genera were represented in more than one module, such as *Granulosicoccus* in modules 4 and 6.

**Figure 5.**
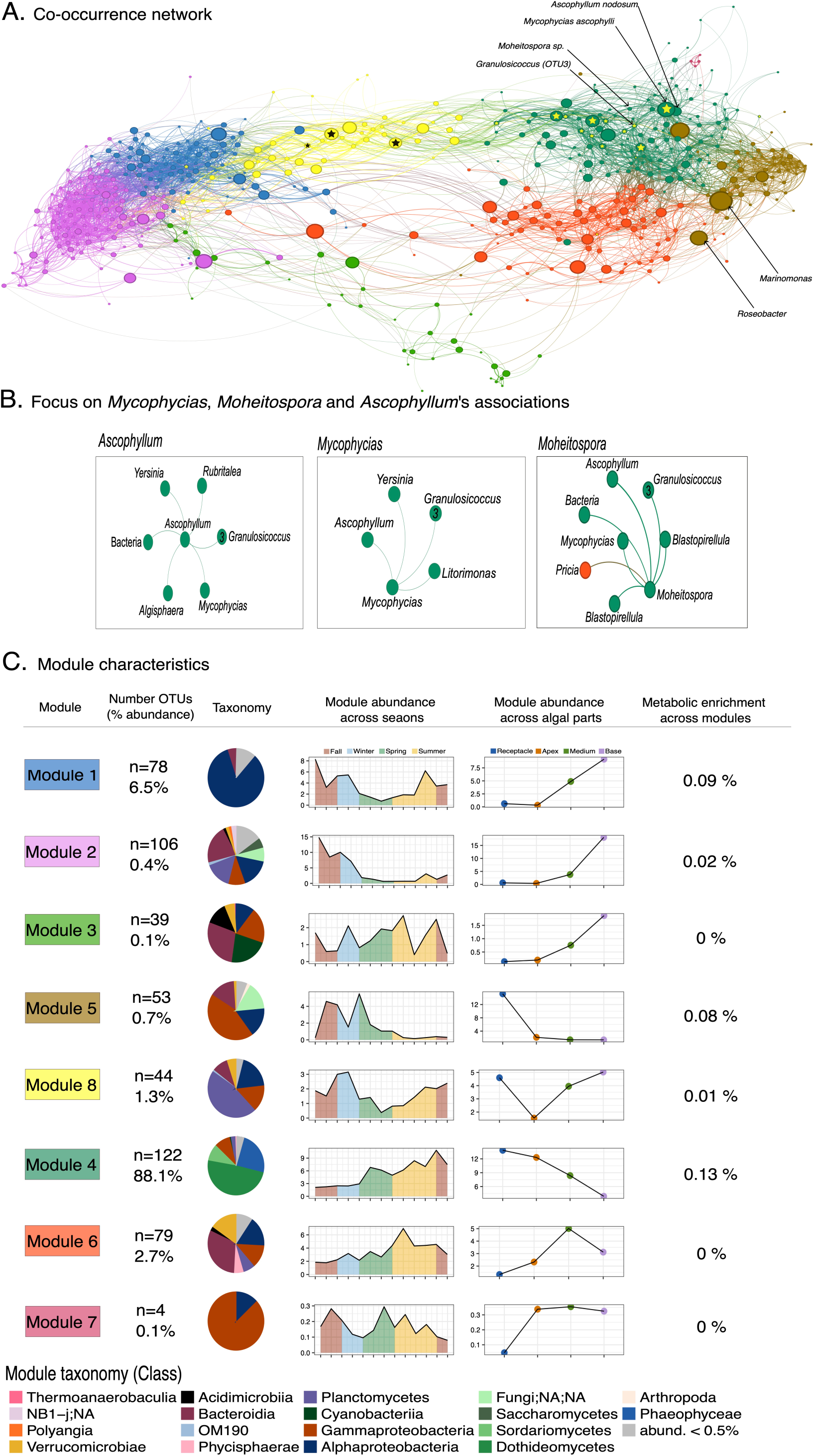
| Co-occurrence analysis of the *Ascophyllum nodosum* microbiota (SSU and ITS). A. Each node represented an OTU (ASVs clustered at 99% identity) while each edge represents a positive correlation obtained from the covariance matrix calculated with SPIEC-EASI. The nodes are colored according to the module they were assigned to, and node size illustrates the betweenness centrality. The network was represented using Gephi and the Force Atlas layout algorithm. Yellow and black stars represent OTUs found in the core microbiome. Node with indicated names seem important as high betweenness centrality or directly in association with *Ascophyllum.* B. Specific interactions with *Ascophyllum nodosum*, *Mycophycias ascophylli,* and *Moheitospora sp.* are illustrated by selecting edges with weight < 0.2. C. Presentation of the 8 modules detected with the Leiden algorithm, ordered by the trend observed in algal part abundance, with the number of OTUs (n) and the relative abundance of each module. The taxonomic composition of the module at class level is illustrated by the pie charts, and the relative microbial abundance across seasons and algal parts is shown in the plots to the right. The Y axis represent the average ASVs abundance found in each algal part or across each month (X axis), where each sample was normalized by the total sum of the ASV in all samples, so each ASV has the same weight. The functional predictions based on the taxonomy were estimated with EsMeCaTa and the percentage of enriched functions in each module is given in the last column.

By adding the measure of the betweenness centrality, we identified key nodes that have an impact on the network structure. The larger the node, the more important the OTUs is likely to be within the interactions. Few larger nodes were observed in the modules 2 and 3, whereas the largest were found in the module 5, associated with the *Maribacter* and *Roseobacter* genera. *Mycophycias ascophylli* and other members of the core microbiome were amongst the largest nodes in module 4 (Figure 6A and Supplementary table 1). The strongest associations of the host *Ascophyllum* were with the OTU_3 of *Granulosicoccus*, *Mycophycias*, *Rubritalea*, *Yersinia*, *Algisphaera*, and one unclassified bacterium (Figure 5B).

**Figure 6.**
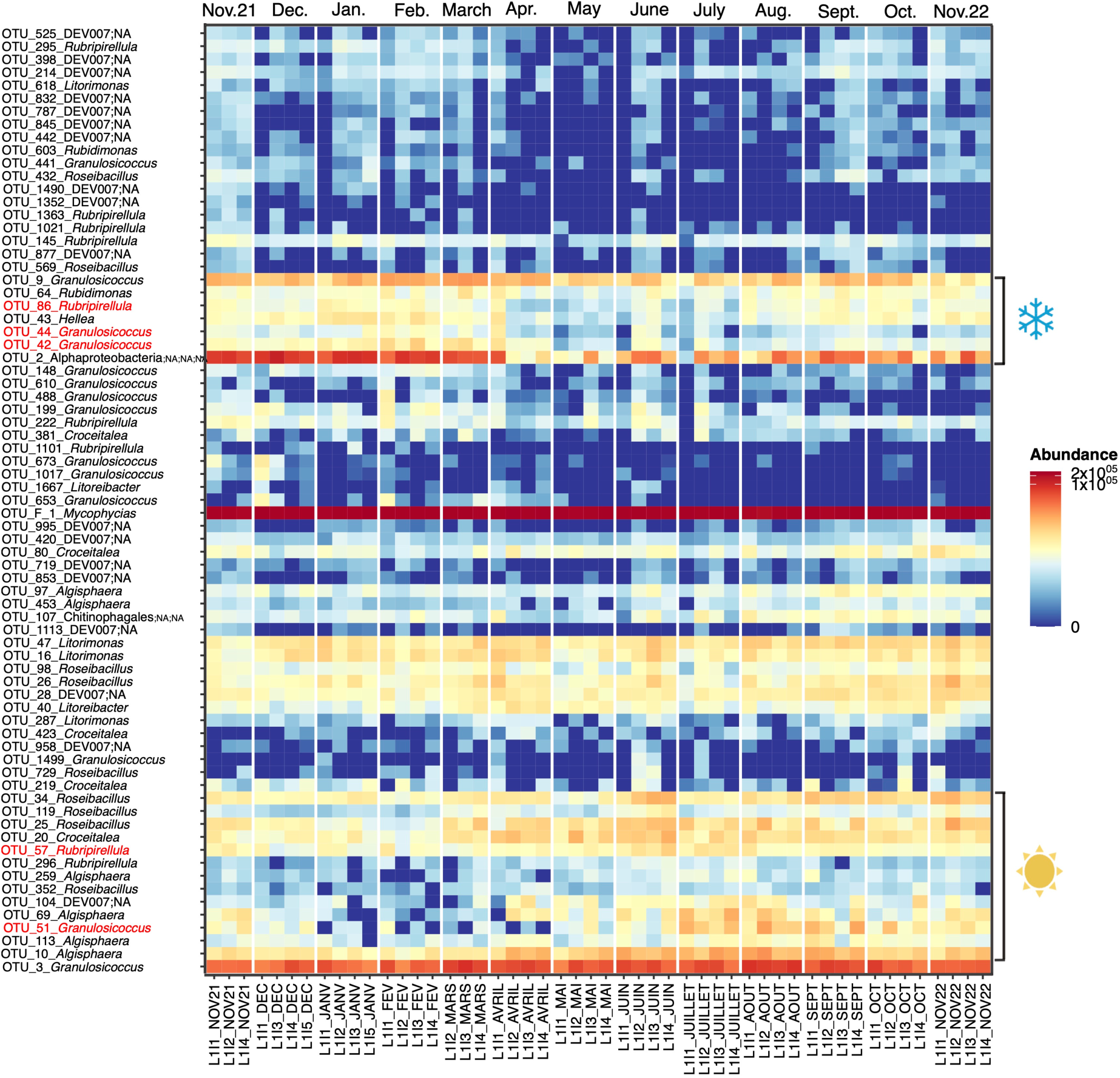
| The heatmap shows the variation of genera found in the core microbiome of *Ascophyllum nodosum* across the monthly survey. The OTUs come from the co-occurrence network ordered by a Pearson correlation. The stronger the red, the more abundant is the OTUs (log10). OTU names in red colors indicate genera with different OTUs that were characteristic for warmer and colder months.

We also investigated the correlation between module distribution and metadata such as seasons and algal tissues (Figure 5C). Microbes associated with modules 1, 2, and 3 were linked to the basal part of the alga whereas module 5, observed at the opposite in the network, was linked to the receptacles from November 2021 to March 2022. Modules 4, 6, and 7 were present all the year. We further investigated the metabolic enrichments based on the taxonomic composition of each module to understand why these microorganisms co-varied together. Among 23,585 predicted functional annotations (GO terms or EC numbers), few were enriched in specific modules (maximum 0.13% for module 1), for example module 5, which was enriched in the siderophore metabolic process (Supplementary Figure 6 and Supplementary Table 2).

### 7. Seasonal dynamics of OTUs within the stable genera of the core microbiome

Prompted by the observation of OTUs corresponding to the same core genera in different modules, we also performed detailed investigation of seasonal variations among the OTUs belonging to genera of the core microbiome. Three groups of OTUs emerged (Figure 6): one exhibiting a stable pattern over the year, one most abundant in colder months between November 2021 and March 2022, and another one mostly found between April and November 2022. Our analyses confirm that some of core genera had OTUs in the different groups. Notably, the *Granulosicoccus* OTU_3 was highly stable, while OTUs 42 and 44 were most abundant in winter, and OTU_51 was most abundant in summer. Similarly, for *Rubripirellula* we observed OTU_86 with a pattern of highest abundance in winter, OTU_57 was dominant in summer, and OTU_145 was relatively stable. This variability suggests the succession of different ecotypes within the core genera.

## Discussion

### 1. Bacterial vs fungal communities: complex and dynamic vs simple and stable

In this study, we explored the *Ascophyllum* microbiome from multiple angles, examining its composition under natural conditions across nearby sampling sites, different parts of the algal thallus, and season over a one-year *in situ* survey. Although the eukaryotic component was observed in *A. nodosum*, the bacteria dominated (98% of the symbiont SSU reads, but extraction biases and SSU copy number differences are likely to impact this figure). Archaeal sequences (<0.001%) were rare and mostly chimeras, suggesting that archaea are absent in *A. nodosum,* unlike in the brown alga *Laminaria rodriguezii* [47]. This supports the idea that bacteria are key microbial symbionts [2].

*A. nodosum* harbors a complex and dynamic bacterial community, in contrast to the stable and simple fungal communities. The bacterial communities comprised 7 different classes with multiple genera, which showed diverse dynamics between months. Microbial composition varied with season and tissues types, aligning with previous studies on macroalgal-associated bacteria [27, 48–51]. Microbial dynamics across tissue types could be related to the physiology and the age of the alga. Given *Ascophyllum*’s apical growth [10], the apex likely host an establishing microbiome, while the basal region likely harbors a more mature microbiome.

In contrast, eukaryotic analysis revealed the predominance of only two fungi: *Mycophycias ascophylli,* and *Moheitospora* sp. (recently renamed *Juncigena* sp.), belonging to Dothideomycetes and Sordariomycetes, respectively. Our study confirms that *Moheitospora* sp. is a central component of the fungal community in *A. nodosum* as shown previously by Vallet *et al.* [19] through culture. Both of these two fungi could contribute to the so-called “lichenization” of *A. nodosum* [14]. Unlike *A. nodosum*, other brown and red macroalgae usually exhibit higher fungal diversity [24, 52, 53]. This suggests a regulation of fungal colonization originating either from the host, the bacteria, or the fungi. Putative antifungal molecules could be synthetized by fungi, as reported in the marine fungi *Aspergillus* sp., and *Penicillium* sp. [54–56]. Bacteria may also control fungal presence, since antibiotic treatment of *A. nodosum* result in *M. ascophylli* hyphae penetrating algal cells and degrading the host [22]. Genome sequencing of these symbionts could provide further indication regarding these hypotheses. The role of *Moheitospora* sp. could also be tested experimentally using its available isolate [19] as shown for *M. ascophylli* [16]. Another perspective is to look at the viral partners as they remain largely understudied, with only a few reports detailing the viral communities associated with seaweeds [57–61]. This lack of knowledge of viruses may limit our capacity to understand the full spectrum of processes occurring within the *A. nodosum* holobiont.

### 2. Host normalization highlights epidermal shedding as a potential driver of bacterial dynamics over the year

In general, metabarcoding analysis only provides relative quantification of the microbiome, although qPCR has been used to mitigate this restriction [27, 62]. Here, we circumvent this limitation without further experimentation, by leveraging our access to the *A. nodosum* 18S rRNA gene allowing us to calculate the proportion of microbial to host reads. We also evaluated the plastid 16S reads for this normalization, but found them to differ up to 5-fold compared the nuclear marker, demonstrating that the two genes cannot be used interchangeably for normalization. Indeed, at least in Archaeplastida, chloroplast abundance does not only vary with tissue type but may be actively regulated by plants depending on the environment [63], making the 18S rRNA gene the likely better choice for normalization.

Based on the proportion of microbial to nuclear host reads, we conclude that the abundance of *A. nodosum*-associated bacteria relative to the host varies over the year. This is globally in line with a previous study by Brunet et al. [27], but our analysis also highlights a drastic drop in bacterial abundance in May, possibly due to epidermal shedding. This process was first observed in 1981 where small pieces of *A. nodosum* were detached occasionally from the thallus surface [64] and considered an adaptation for long-lived macroalgae to control epiphytes [65, 66]. The phenomenon was observed [67] and quantified weekly, revealing regular fluctuations of the surface microbiome throughout the warmer months of the year [68]. However, we observe that the reduction in the ratio of microbial to host reads did not concern all microbes. A large proportion of the unassigned Alphaproteobacterial ASV and other bacteria seem to be impacted by the process, while, e.g., one *Granulosicoccus* OTUs or the two main fungi remained abundant over the year.

In the context of shedding, this would mean that these consistently abundant organisms are either fast colonizers rapidly taking hold on the free surface, or that these microbes also penetrate into deeper cell layers and are less affected by shedding. The latter is likely true at least for the most abundant of the two fungi, *M. ascophylli*, which has even been found intracellularly [8] or in the intercellular spaces embedded in the thick cell walls of the medulla and cortex tissues [69]. Regardless of the reasons driving these different patterns, our observations suggest that even within a genus, distribution patterns and thus the underlying ecological niches and strategies of different OTUs differ. Fine taxonomic distinction will therefore be crucial in future investigation of these taxa.

### 3. Stability revealed: 22 key ASVs in the core microbiome of *A. nodosum*

Identifying core microbiomes has become widespread to identify potential key actors in host biology. In our study, the metric for quantifying the core microbiome is 100% occupancy, meaning that microbial taxa need to be present in every sample to be considered in the core. While this is a very stringent measure [70], the overall high stability of the *A. nodosum* microbiome in combination with the high sequencing coverage make this a viable choice for our dataset.

Among the 22 core amplicon sequence variants (ASVs) present in all *Ascophyllum* individuals, several genera, in particular *Granulosicoccus*, *Litorimonas, Hellea, Blastopirellula,* and *Roseibacillus* are generalists that have been observed on a diverse array of brown, green, and red macroalgae [27, 50, 51, 71–78]. The ubiquity of the genus *Granulosicoccus* in several brown algae suggests that it is especially well-suited for life as a symbiont. Metagenome-assembled genomes (MAG) of *Granulosicoccus* were annotated in the bull kelp *Nereocystis luetkeana*, and harbored genes for synthesizing cobalamin possibly providing this and other vitamins to the algal host [79]. However, evidence of metabolic exchanges between *Granulosicoccus* and the algal host has not yet been completed.

Our core microbiome analyses also consolidate the tight association of *M. ascophylli* (Dothideomycetes;NA) with the *Ascophyllum*, as all samples contained the fungus in high abundance. The symbiotic association between *Ascophyllum* and this fungus has already been described and reviewed [8, 14, 53, 80–83]. Previous light microscopy analyses have shown *M. ascophylli* to be present at the highest concentration in receptacles [15], which is not consistent with our findings, but this likely point to a limitation of normalization method – indeed the male receptacles produce very high numbers of sperm cells each with a complete copy of the host genome thus potentially biasing the number of host reads used as a reference in this case. Regardless of these differences, the constant presence of the fungus is consistent with its supposed role in host development, fitness, and protection against biotic and abiotic stresses [16, 84].

### 4. Co-occurrence network reveals microbial contributions to energy synthesis in receptacles

Our co-occurrence networks analysis revealed that *A. nodosum* was not the central node in the network, and the identified central nodes are not those in association with the host, except for *M. ascophylli*. Very few nodes directly in association with the seaweed. This suggests that *Ascophyllum* may be a surface for few key microbial colonizers (whether endophytic or epiphytic), as *Mycophycias* and *Granulosicoccus,* which may be the base for the biofilm formation. These analyses also highlight module 4, which appears to be key module, as it is the most abundant module in terms of read coverage and contains many species present in the core microbiome. Module 4 could be linked to early-colonizing or intracellular microorganisms, such as those not responding to shedding phenomenon, and may include functions that contribute to the holobiont’s overall stability, while e.g. module 5 may be linked to reproduction.

Because modules regroup highly connected OTUs that share the same environment and potentially exhibit coherent ecological strategies [85], we expected to find the modules to be enriched in functions related to niche adaptation. One example of such adaptations concerned the functioning of receptacles. In the reproductive period, energy should be allocated to reproduction and not to vegetative growth [13]. Iron, essential for ATP production in photosynthesis and respiration, is present in low concentration and poor solubility in seawater [86, 87]. To facilitate iron acquisition, many bacteria synthesize siderophores [88]. Functions present in the module 5 were enriched in siderophore metabolic processes (GO: 0009237, Supplementary figure 6), with *Roseobacter* and *Marinomonas,* two bacteria likely possessing genes involved in siderophore production [89] (Supplementary Table 1), occupying structuring positions in the receptacles. As these enriched metabolic functions are merely predictions based on the microbial taxonomy, they need to be confirmed by complementary approaches, such as metagenomic analyses. Overall, our analyses highlight that very few metabolic functions are enriched across modules, underscoring the functional stability of the holobiont.

## Conclusion and perspectives

Our study provides a comprehensive overview of the microbial communities associated with *Ascophyllum nodosum,* highlighting the complexity of its holobiont and the dynamic interactions between its bacterial and fungal components. Using metabarcoding analysis with universal primers and using the host 18S for normalization, we found that *Ascophyllum* harbors a predominantly bacterial microbiota, with no detectable archaea. Community composition and structure vary across algal tissues and seasons, including a notable drop in bacterial abundance in May, potentially linked to epidermal shedding. In contrast, the fungal communities in *Ascophyllum* appear stable and simple, primarily consisting of *M. ascophylli* and *Moheitospora* sp., both of which appear to maintain a close association with the host throughout the year. Our identification of a core microbiome with 22 ASVs suggests the importance of certain microbial taxa in maintaining host biology, such as the genus *Granulosicoccus*. While the latter genus demonstrates significant stability, our findings also reveal that within this genus different OTUs are subject to seasonal variation, highlighting its intraspecific diversity and adaptability. The co-occurrence network and its module-based analyses add another layer of complexity to our understanding, showing how microbial groups interact in response to ecological niches, such as parts of the thallus or varying seasonal conditions. Further investigation using metagenomic and functional approaches will be key to uncovering the adaptations and the roles of key microorganisms and understanding their contributions to the overall health and stability of the *A. nodosum* holobiont.

## Acknowledgments

We would like to thank Cécile le Guillard (Plant Nutrition R&D Department, Centre Mondial de l’Innovation of Groupe Roullier, 35400 Saint-Malo, France), Soizic Prado and Gautier Demoulinger (Molecules of Communication and Adaptation of Microorganisms, UMR 7245, MNHN, CNRS, 75000 Paris, France and Plant Nutrition R&D Department, Centre Mondial de l’Innovation of Groupe Roullier, 35400 Saint-Malo, France) for discussions and help during sampling. We would like to thank Nicolas Henry (ABiMS, FR2424, Station Biologique de Roscoff, Sorbonne Université, CNRS, 29680 Roscoff, France) for discussions on universal primers and Loïs Maignien (Biology and Ecology of Deep-Sea Ecosystems, IUEM, 29280 Plouzané, France) for his help with co-occurrence network. We thank Gaëtan Burgaud (LUBEM, Université de Bretagne Occidentale, 29280 Plouzané, France) for his help for ITS primer choices and taxonomic assignations and for his advices during manuscript reviewing. We would like to thank Pauline Hamon-Guiraud (Dyliss, Inria, UMR 6074, CNRS, University of Rennes, Rennes, France) for help in EsMeCaTa tool. We thank François Thomas (Integrative Biology of Marine Models, UMR 8227, Station Biologique de Roscoff, Sorbonne Université, CNRS, 29680 Roscoff, France) for discussions around microorganisms and for reviewing this manuscript. We also thank for their support, the Genomer platform and the Roscoff Bioinformatics platform, ABIMS (http://abims.sb-roscoff.fr), respectively part of EMBRC-France (financially supported by the Investments of the Future program, ANR-10-INSB-02) and of the Institut Français de Bioinformatique (ANR-11-INBS-0013), and of the BioGenouest network. We are most grateful to the Genomics Core Facility GenoA, member of Biogenouest and France Genomique and to the Bioinformatics Core Facility BiRD, member of Biogenouest and Institut Français de Bioinformatique (IFB) (ANR-11-INBS-0013) for the use of their resources and their technical support.

## Author contributions

CR, CL and SD designed and conceptualized the research. CR collected samples, performed laboratory work, analyzed data and wrote the manuscript. GT and EL performed sequencing preparations. SB and AB implemented the EsMeCaTa tool and contributed to the functional analyses. SR helped for sampling and DNA extractions. PP discussed around algal physiology, provided funding and reviewed the manuscript. CL and SD collected samples, supervised the research, reviewed the manuscript and provided funding. All authors have read and approved the manuscript.

## Data availability

The raw sequence reads and metadata are available in the European Nucleotide Archive (ENA) database under BioProject PRJEB85975 and sample accession numbers ERS23809686-ERS23809790. ASV table with final abundances, taxonomy, and sequences are available in supplementary tables. Scripts are available on GitHub (https://github.com/rssco/novaseq_ascophyllum).

## Supplementary figures

**Supplementary Figure 1.**
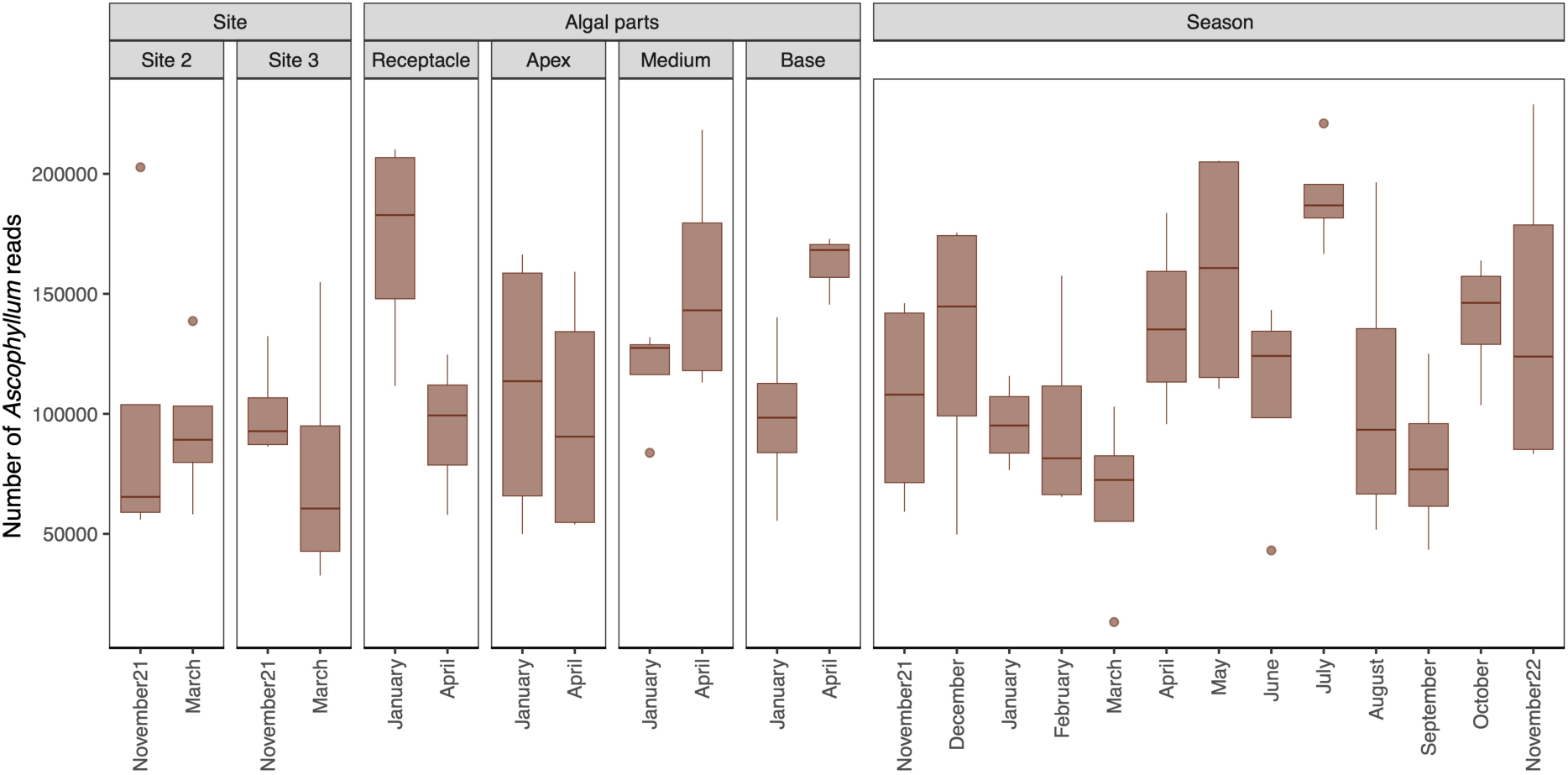
| Distribution of *A. nodosum* read abundance across the three datasets

**Supplementary figure 2.**
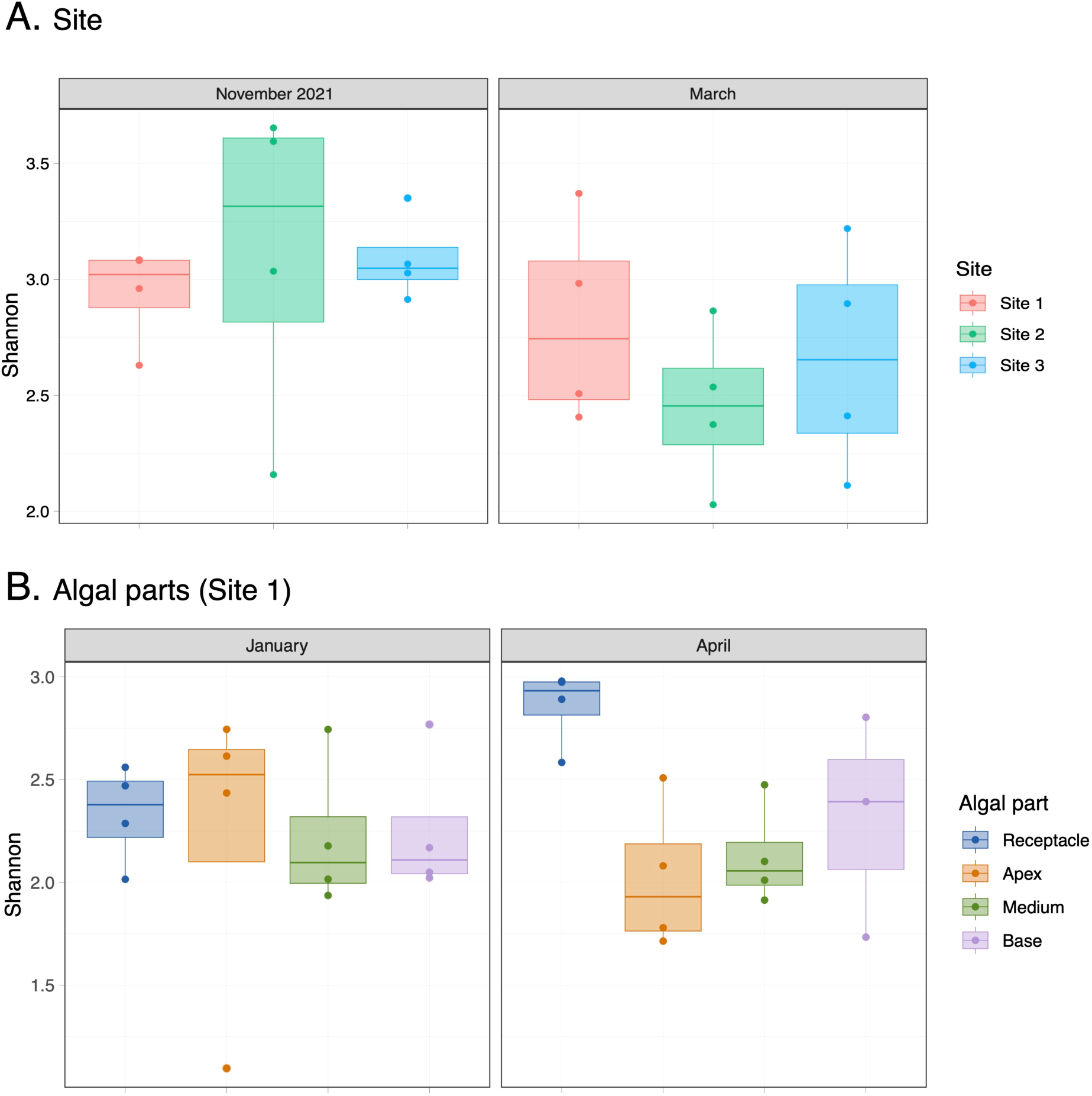
| Alpha diversity for SSU (Shannon index) of the sites (A) and the algal part datasets (B). No significant differences are observed according to an ANOVA test.

**Supplementary figure 3.**
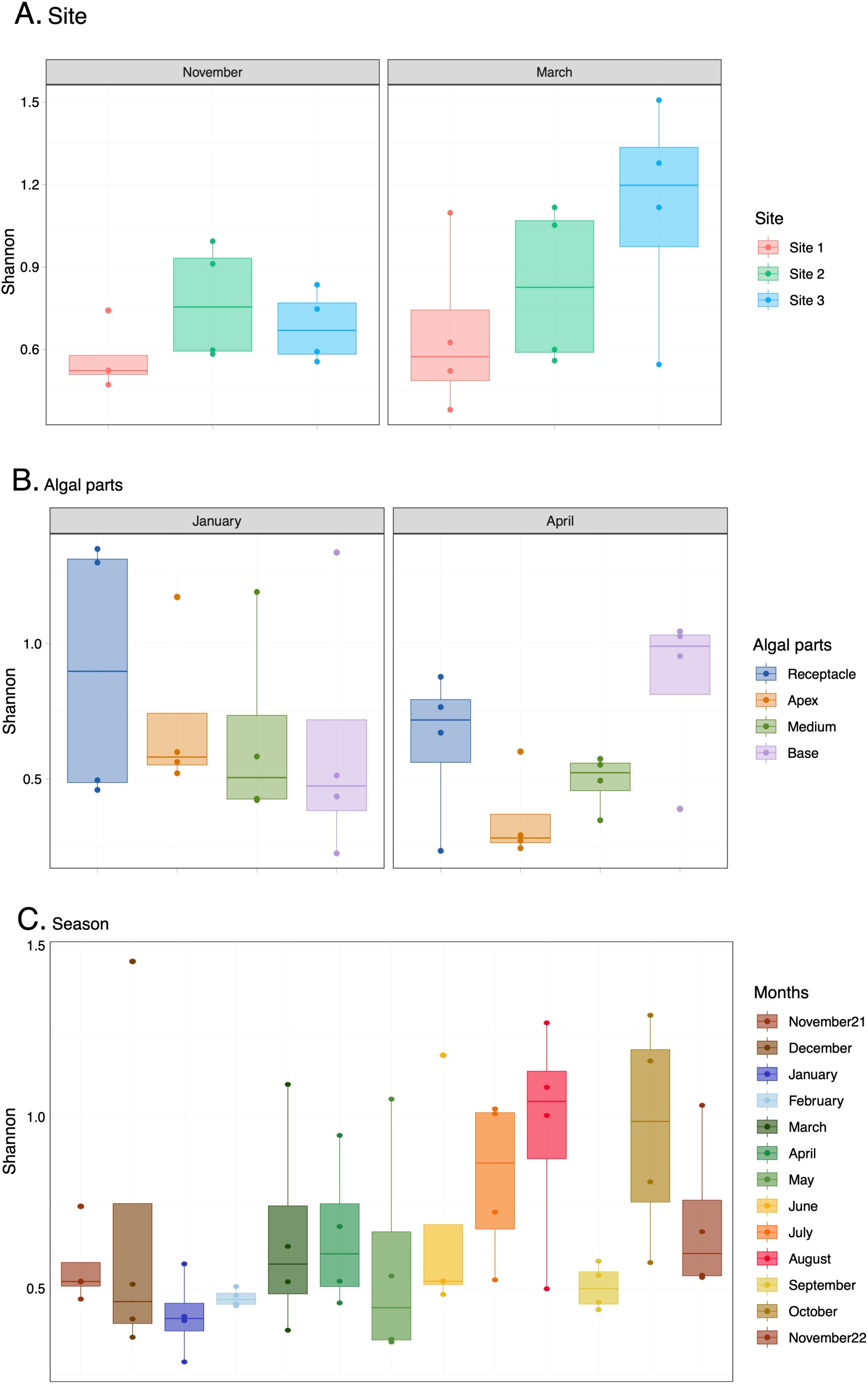
| Alpha diversity for ITS (Shannon index) of the sites (A), the algal part (B) and season (C) datasets. No significant differences are observed according to an ANOVA test.

**Supplementary figure 4.**
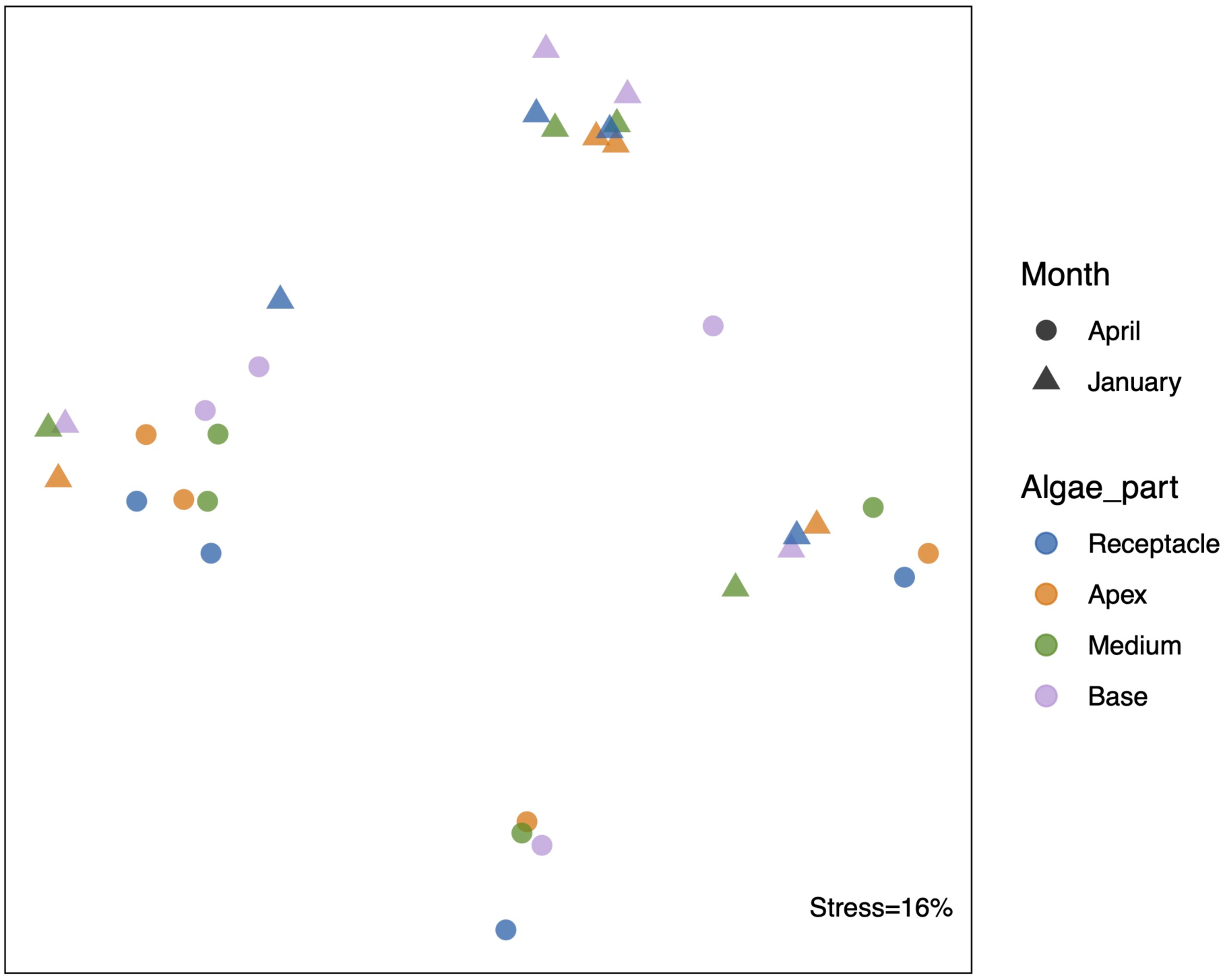
| Beta diversity of the fungal communities across the four algal parts, showing significant results between month (p-value=0.010). Statistical analyses were based on a PERMANOVA test.

**Supplementary figure 5.**
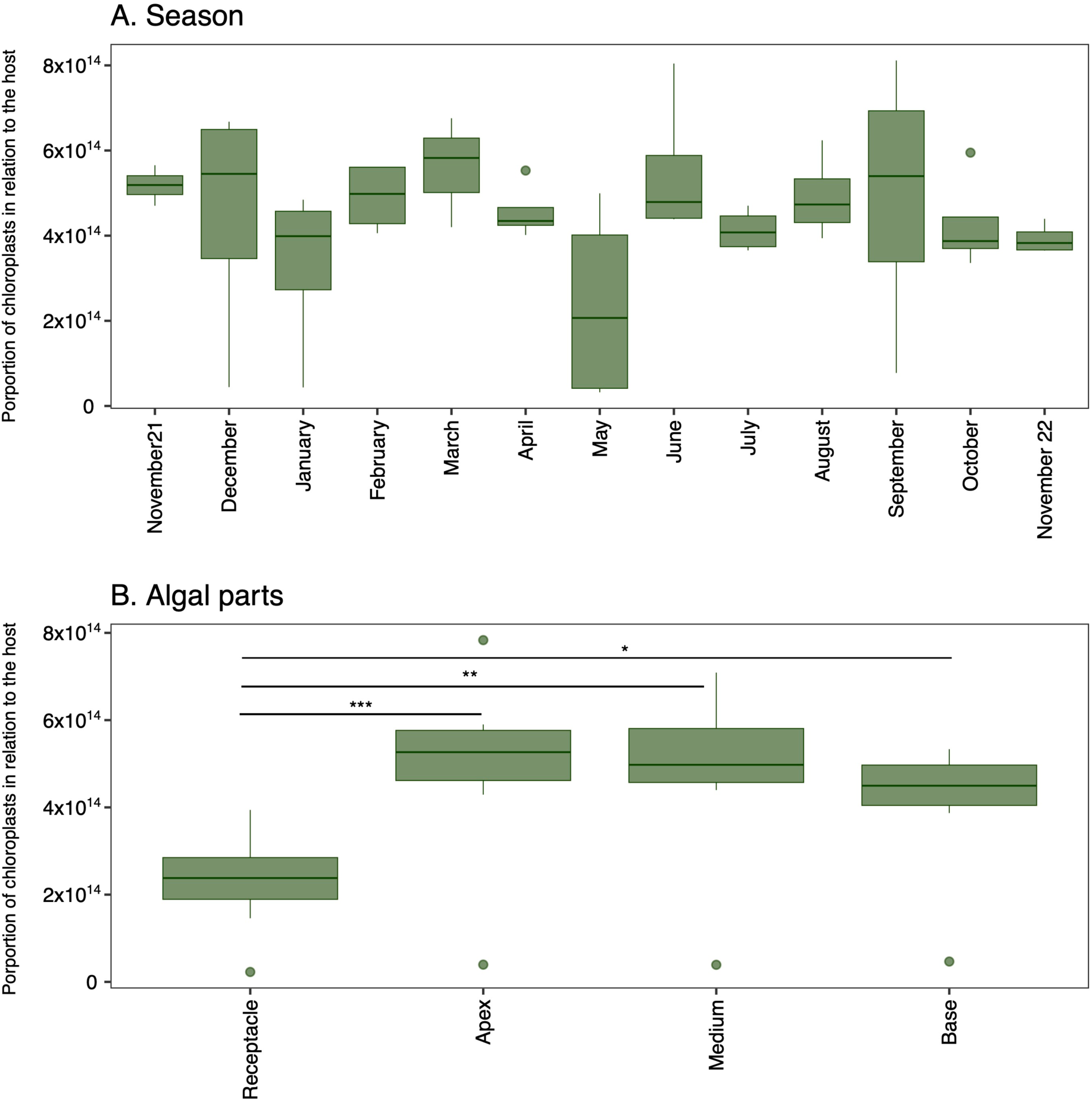
| Following the ratio of plastid read to host reads over each month of the year (A) and across algal thallus parts (B). Significant differences according to an ANOVA (non-significant for season, p = 0.0005 for algal parts) test with a Tukey post-hoc test are indicated by different *

**Supplementary figure 6.**
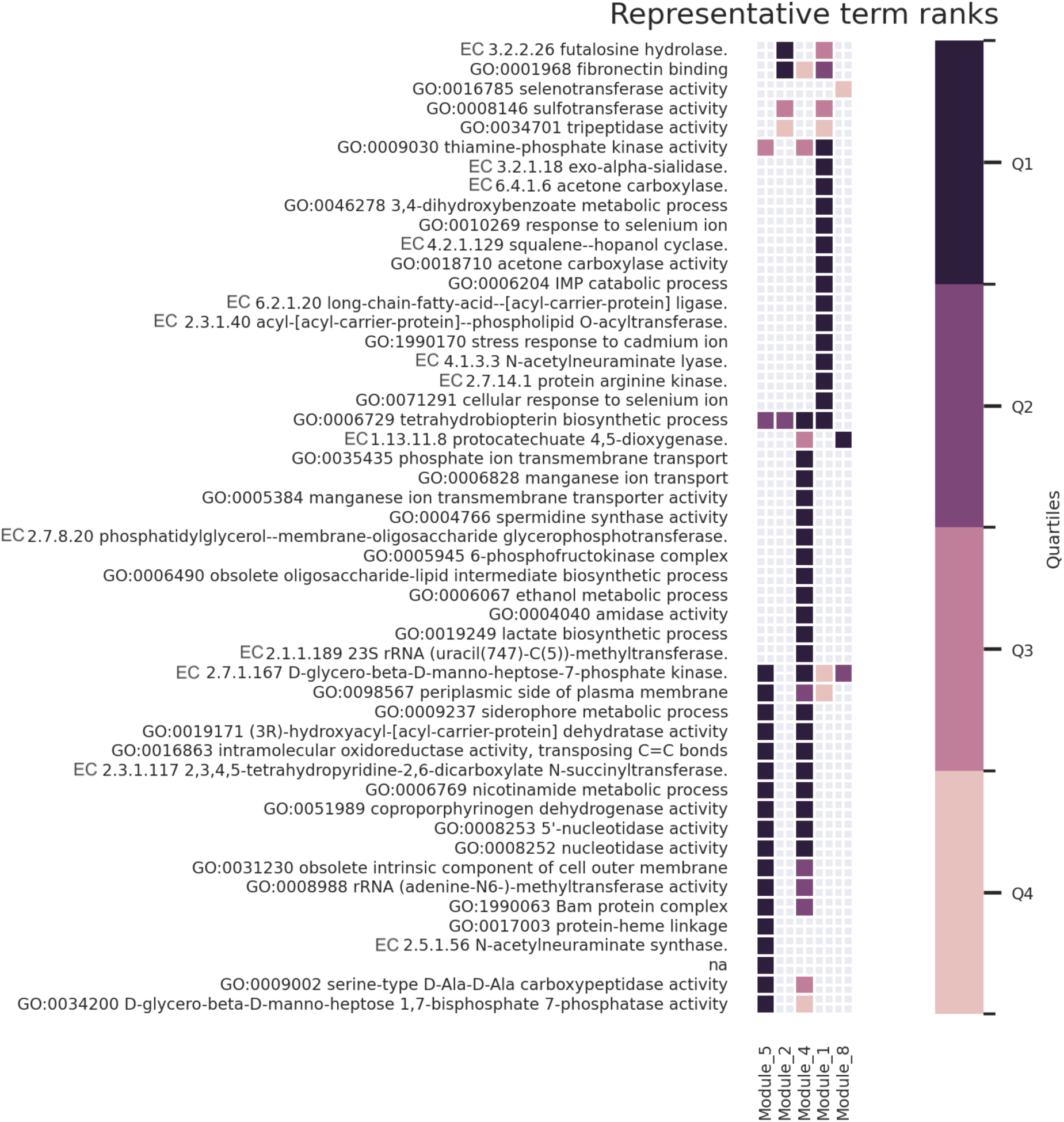
| Metabolic enrichment of GO terms and EC number predictions of 5 modules present in the co-occurrence network. Annotation predictions were based on taxonomic affiliation with EsMeCaTa tool. Module 3, 6 and 7 are not enriched and not represented here. Metabolites in Q1 corresponds to high enrichment.

## Supplementary tables

Supplementary table 1 | Betweenness centrality of each ASV present in the co-occurrence network and their attributed module. The abundance of each ASV is also indicated.

Supplementary table 2 | Amplicon sequences variants module 5 which are estimate to have gene in siderophore metabolic process (GO:0009237).

Supplementary table 3 | ASV table with abundance, taxonomy and references for SSU dataset

Supplementary table 4 | OTU table with abundance, taxonomy and references for ITS dataset.

